# *Clostridium scindens* colonization of gnotobiotic mice promotes a chronic unresolving infection with *Clostridioides difficile*

**DOI:** 10.1101/2022.06.12.495821

**Authors:** M Graham, N DiBenedetto, ML Delaney, R Lavin, A Pavao, V Yeliseyev, L Bry

**Affiliations:** Massachusetts Host-Microbiome Center, Dept. Pathology, Brigham & Women’s Hospital, Harvard Medical School. Boston, MA 02115; Department of Molecular Biology and Microbiology, Tufts University School of Medicine, Boston, MA, 02111; Clinical Microbiology Laboratory, Dept. Pathology, Brigham & Women’s Hospital, Harvard Medical School. Boston, MA 02115

## Abstract

The commensal *Clostridium scindens* has been regarded as a promising bacteriotherapeutic against *Clostridioides difficile* infection due to its ability to consume host factors that can promote *C. difficile* growth, and its production of the antimicrobial compound 1-acetyl-β-carboline. We investigated *C. scindens’* protective effects against *C. difficile* using defined colonization studies in gnotobiotic mice. Mice infected with *C. difficile* develop lethal infection within 48 hours. In contrast, 88% of mice pre-colonized with *C. scindens* survived acute infection with delayed *C. difficile* colonization, lower biomass, and toxin B levels at 24 hours after infection. However, two weeks post-challenge, surviving mice showed comparable levels of cecal *C. difficile* vegetative and spore biomass and toxin B, as seen during acute infection. After two weeks, co-colonized mice exhibited mucosal colonic hyperplasia with focal pseudomembranes, modeling a chronic and recurrent infection state. Our findings illustrate how the commensal microbiota can modulate host and pathogen interactions leading to chonic *C. difficile* carriage and infection.

## Introduction

*Clostridioides difficile* is a common nosocomial infection that causes substantial morbidity and mortality (*1, 2*). *C. difficile* causes over 450,000 infections and 20,000 deaths annually in the US and is a major public health threat worldwide (*3, 4*). *C. difficile* infection is associated with antibiotic use (*5*), which perturbs protective gut microbiota, allowing *C. difficile* to proliferate and cause infection (*6*). Antibiotics targeting vegetative *C. difficile* cells are routinely used to treat infection, but up to 35% of patients who respond initially suffer recurrent infections, likely in part due to the persistence of *C. difficile* spores (*7*). Understanding the factors that contribute to recurrent infections has been challenging, as well as how improved use of antibiotics and or fecal microbiota transplants (FMT) or targeted bacteriotherapeutics can prevent their occurrence (*8*).

The commensal *Clostridium scindens* has been associated with resistance to *C. difficile* infection clinically and in animal models. *C. scindens* is a cluster XIVa *Clostridia* that ferments common hexoses and pentoses (*9*). It also possesses systems to ferment proline and glycine via reductive Stickland pathways (*10–12*). *C. scindens’* proposed mechanisms of action against *C. difficile* have included the conversion of primary to secondary bile acids via 7α-hydroxysteroid dehydrogenase activity to inhibit *C. difficile* spore germination (*13*), and production of the antimicrobial compound 1-acetyl-β-carboline, which inhibits *C. difficile* vegetative growth *in vitro* (*14*). However, studies by our group and others have suggested that metabolic competition commensal *Clostridia* contributes to their beneficial effects *in vivo* (*12, 15, 16*).

We evaluated *C. scindens’ in vivo* interactions with toxigenic *C. difficile* using a defined-association infection model in gnotobiotic mice (*12*). We reveal the commensal’s effects on disease phenotypes including the suprising finding of promoting a chronic and unresolving infection. The findings provide new insights into commensal drivers of acute and chronic *C. difficile* infection, including in an otherwise healthy and immunocompetent host.

## Results

*C. difficile* infection caused lethal disease in 6-week-old germ-free mice by 48 h of infection (Figures 1A and 1B). Mice displayed symptoms at 20 h including diarrhea and deteriorating body condition. Mice exhibited worsened disease, epithelial damage, and transmural neutrophilic infiltrates in the colon at 24 h post-infection (Figure 1F).

**Figure 1.**
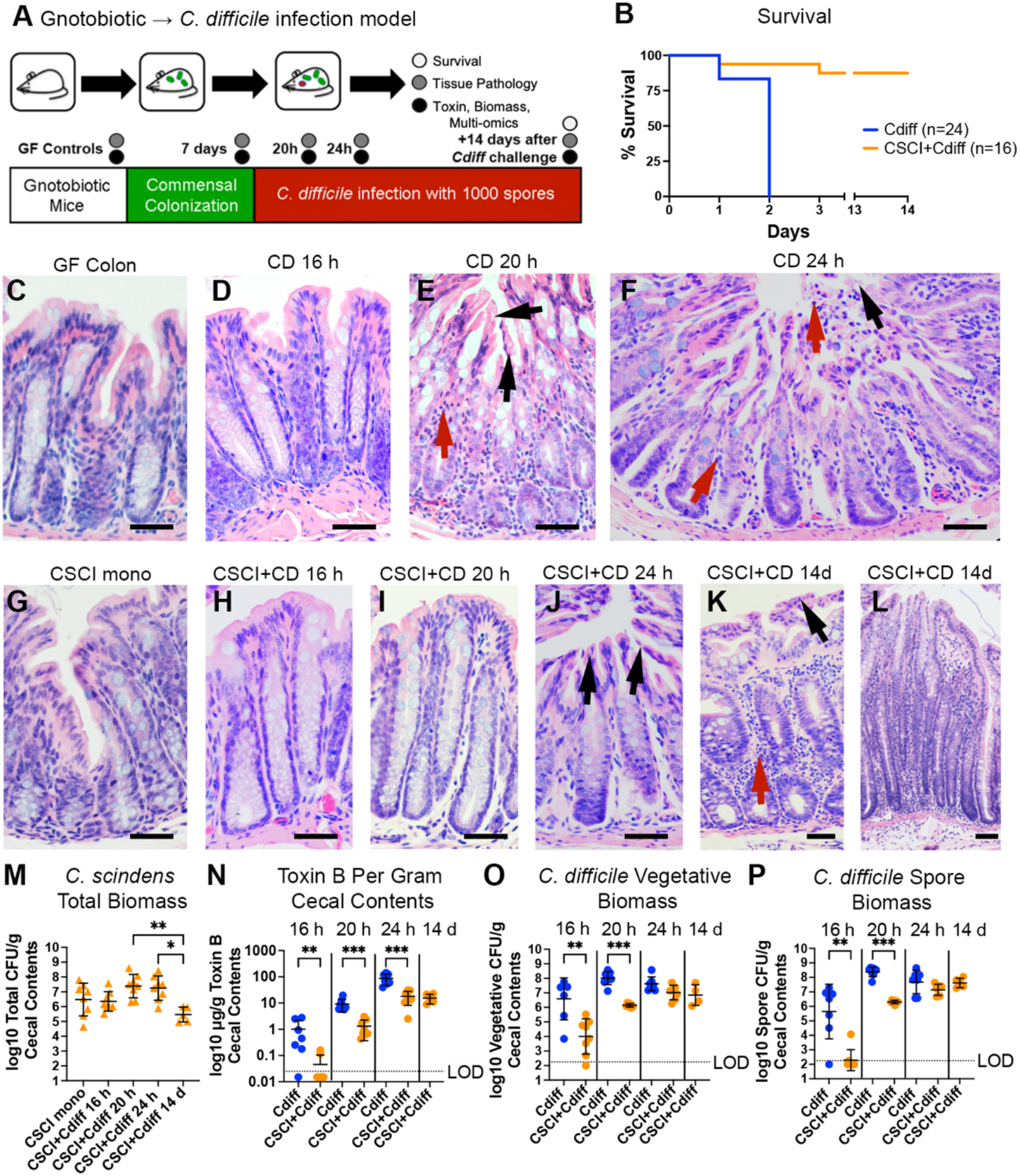
*C. scindens* protects germ-free mice from acute, lethal *C. difficile* infection and promotes chronic, unresolving disease. **A)** Experimental overview. **B)** Survival curves. **C-K)** Hematoxylin and eosin (H&E)-stained colon sections. Magnification: 200X, scale bar 50 μm (C-K) and 100X, scale bar 50 μm (L). **C)** Normal germ-free mucosa. **D)** *C. difficile*-infected mice at 16 h of infection showing normal epithelium. **E)** *C. difficile*-infected mice at 20 h of infection showing vacuolation and epithelial stranding (black arrows), and neutrophilic infiltrates (red arrow). **F)** *C. difficile*-infected mice at 24 h showing surface epithelial loss (black arrow) and transmural neutrophilic infiltrates entering the lumen (red arrow). **G)** *C. scindens* monocolonized mice. **H)** *C. scindens*- and *C. difficile*-infected mice at 16 h. **I)** *C. scindens*- and *C. difficile*-infected mice at 20 h. **J)** *C. scindens*- and *C. difficile*-infected mice at 24 h showing epithelial stranding and vacuolation (black arrows), but little inflammation. **K, L)** *C. scindens*- and *C. difficile*-infected mice at 14 d after infection showing neutrophilic infiltrates (red arrow), formation of focal pseudomembranes (black arrow), and (**K**) marked mucosal hyperplasia (**L**). **M)** Log_10_ *C. scindens* total biomass CFU/g cecal contents. Bars show mean and standard deviation. Kruskal-Wallis significance values: * 0.01 < p ≤ 0.05; ** 0.001 < p ≤ 0.01. **N)** Log_10_ extracellular cecal Toxin B (μg/g) cecal contents. Bars show mean and standard deviation. Limit of detection denoted by LOD. Mann-Whitney significance values: ** 0.001 < p ≤ 0.01; *** 0.0001 < p ≤ 0.001. **O)** Log_10_ *C. difficile* vegetative CFU/g cecal contents. Bars show mean and standard deviation. **P)** Log_10_ *C. difficile* spore CFU/g cecal contents. Bars show mean and standard deviation.

Mice pre-colonized with *C. scindens* prior to *C. difficile* challenge survived infection (Figure 1B; p<0.0001) with milder symptoms and reduced colonic damage. Fourteen days post-infection, animals showed further colonic damage and inflammation with the formation of focal pseudomembranes and hyperplastic mucosa (Figure 1 K-L).

*C. difficile* monocolonized animals had higher cecal vegetative and spore biomass as well as higher extracellular toxin B levels initially. By 24 h post-infection, *C. scindens* pre-colonized animals demonstrated similar biomass to *C. difficile* monocolonized animals, but with 5-fold lower toxin B levels. After 14 days, vegetative and spore biomass, and toxin B levels remained at acute levels in surviving *C. scindens* pre-colonized mice, indicating long-term infection.

*C. scindens* biomass remained at comparable levels in mono-colonized mice and during the acute phase of infection after *C. difficile* challenge. However, after 14 days, *C. scindens* biomass decreased relative to early *C. difficile* infection (Figure 1 M).

## Discussion

Using a gnotobiotic model of *C. difficile* infection, we demonstrate the partial protection *C. scindens* confers in mice infected with *C. difficile*. While *C. scindens* reduced acute lethality from infection, it permitted long-term pathogen carriage and chronic disease, modeling a chronic, recurrent infection.

*C. difficile’s* capacity to colonize and the duration over which it produces toxin in the gut determine its propensity to cause disease. Pre-colonization with *C. scindens* slowed *C. difficile* colonization, delaying the onset of acute infection, reducing toxin load, and protecting against rapidly fatal infection. However, while *C. scindens* supported survival beyond acute, symptomatic infection, it did not prevent chronic toxin production and associated host disease. The maintenance of high *C. difficile* biomass and toxin levels in *C. scindens* pre-colonized mice led to chronic mucosal damage and a hyperplastic mucosal response. The chronic nature of the colonic inflammation and focal pseudomembranes models that of chronic and recurrent *C. difficile* infections observed clinically (*17*), and provides a tractable mouse model to evaluate these aspects of human disease.

*C. scindens’* ability to convert host primary bile acids to secondary bile acids with anti-germination effects on *C. difficile* spores has previously been proposed as a central mechanism by which it inhibits *C. difficile* growth *in vivo* (*13, 18*). Studies by our group and Aguirre, *et al*. have shown that commensals, particularly Stickland amino acid fermenting species that include *P. bifermentans*, which lacks 7α-hydroxysteroid dehydrogenase activity, as well as *C. hiranonis* and *C. scindens*, promoted survival against lethal *C. difficile* infection in defined-association mouse models (*12, 16*). Detailed studies of *P. bifermentans* and *C. difficile* interactions *in vivo* showed complex molecular actions of protection, including that this Stickland fermenting species competes robustly against *C. difficile* for nutrients, particularly the Stickland-fermentable amino acids proline, glycine, and leucine, in addition to preferred carbohydrates and polyamines. Our establishment of a tractable chronic infection model with *C. scindens* and *C. difficile* provides means to define commensal contributions including metabolic competition, production of antimicrobial factors, and other complex mechanisms of action to further our understanding of drivers for recurrent infections and interventions to better prevent and treat them.

## Methods

### Bacterial strains and culture conditions

*C. difficile* strain ATCC 43255 (ATCC, Manassas, VA) spores were prepared as described (*19*). *C. scindens* strain ATCC 35704 (ATCC, Manassas, VA) cultures were grown in brain heart infusion broth (3.7%) supplemented with hemin (0.0005%) and vitamin K (0.0001%) in a Coy anaerobic chamber (Coy Labs, Grass Lake, MI) in an atmosphere consisting of 80% N_2_, 10% H_2_, 10% CO_2_ at 37°. The concentration was quantitated by serially diluting and counting colonies on Brucella agar (Becton Dickinson, Canaan, CT). The culture was aliquoted into a sterile Hungate tube for transport outside of the chamber.

For *C. difficile* and *C. scindens* biomass, flash-frozen mouse cecal contents were collected into pre-weighed Eppendorf tubes with 1 mL of pre-reduced PBS with 0.05% cysteine (Milllipore-Sigma, St. Louis, MO) as a reducing agent. Tubes were weighed after adding material and moved into the anaerobic chamber for serial dilutions with plating to selective *C. difficile* CHROMID® agar (Biomérieux, Durham, NC) or Brucella agar (Becton Dickinson, Canaan, CT) supplemented with 500uL of PBS with 0.1% taurocholic acid (Millipore-Sigma, St. Louis, MO) for *C. scindens* quantitation. *C. difficile* colonies were counted after 48 hours of incubation, and were identified as large black colonies. *C. scindens* colonies were counted after 48 hours and were identified as small, round colonies.

*C. difficile* spores were isolated by exposing pre-weighed cecal contents in PBS to 50% ethanol for 1 hour followed by serial dilution and plating to CHROMID® agar, as described (*20*). Vegetative cell biomass was calculated by subtracting the spore biomass from the total biomass and normalizing to mass of cecal contents. Data were analyzed in Prism 9.0 (GraphPad, San Diego, CA) for visualization and log-rank tests of significance among groups. A p value ≤ 0.05 was considered significant.

### Mouse Studies

All studies were conducted under an approved institutional IACUC protocol. Equal ratios of 6-7-week-old male and female gnotobiotic Swiss-Webster mice were administered 1×10^8^ of *C. scindens* or sterile vehicle control by oral gavage and allowed to colonize for 7 days prior to *C. difficile* challenge in a positive pressure gnotobiotic isolator (Class Biologically Clean, Madison, WI). Mice were transferred out of the isolator and singly housed in sterile OptiMice ventilated containment cages (Animal Care Systems, Centennial, CO). Fecal pellets were cultured to confirm mono-association with *C. scindens* or maintenance of the germ-free state prior to transfer. Mice were challenged with 1×10^3^ *C. difficile* spores by oral gavage. Progression of disease was assessed via a modified body condition scoring system (*19*). Mice were sacrificed at a BCS of 2- (emaciation, lack of ambulation, anal prolapse) or at the following defined timepoints: 7 days of *C. scindens* colonization or germ-free controls, 16, 20, 24 h, or 14 days post-*C. difficile* challenge. Cecal contents were collected and flash-frozen in liquid nitrogen. The GI tract and internal organs were fixed in zinc-buffered formalin (Z-FIX, Thermo Fisher Scientific, Waltham, MA) for histopathologic assessment.

### Histopathologic Analyses

Formalin-fixed gut segments from germ-free or specifically associated mice were paraffin-embedded and 5 μm sections cut for staining with hematoxylin and eosin (H&E: Thermo Fisher Scientific, Waltham, MA) as described (*21*). Slides were visualized with a Nikon Eclipse E600 microscope (Nikon, Melville, NY) to assess epithelial damage per cellular stranding, vacuolation, mucosal erosions, and tissue edema, and to characterize inflammatory infiltrates.

### Toxin B ELISA

Cecal toxin B levels were quantified as described (*22*). Microtiter plates were coated with 5 μg/mL of anti-TcdB capture antibody (BBI Solutions, Madison, WI). Supernatants of spun cecal contents and standard curve controls of toxin B (ListLabs, Campbell, CA) were assayed in triplicate. After incubation and washing, the plate was incubated with biotinylated anti-toxin B antibody (polyclonal goat-anti-*C. difficile* TcdB; ab252712, Abcam, Cambridge, MA), washed, and incubated with Pierce™ High Sensitivity Streptavidin-HRP conjugate (Thermo Fisher Scientific, Waltham, MA). Signal was developed with TMB substrate (Thermo Fisher Scientific, Waltham, MA) and sulfuric acid and quantified at 450 nm using a BioTek Synergy H1 plate reader (BioTek Instruments Inc., Winoski, VT). Values were analyzed in Prism 9.0 (GraphPad, Sand Diego, CA) to calculate μg of toxin B per gram of cecal contents. Differences among groups were evaluated by non-parametric Mann-Whitney test. A p value ≤ 0.05 was considered significant.

